# Advancing North: White-nosed coati’s *Nasua narica* Range Expansion in Arizona

**DOI:** 10.1101/2022.05.26.493663

**Authors:** Kinley Ragan, Faith M. Walker, Lawrence E. Stevens, Jan Schipper

## Abstract

Over the past century, the white-nosed coati (WNC; *Nasua narica*) has expanded its northernmost range from the United States-Mexico border into northern Arizona. WNC are medium-sized, opportunistic omnivores that often occur in large groups (“bands”) and forage on insects, fruits, and small vertebrates. We compiled data from iNaturalist, published literature, Arizona Game and Fish records, museum collections, personal communications, and our own camera trap photography to chronicle this range expansion. Historical records documented WNC populations in mountainous areas along the US-Mexican border but rarely north of Tucson, AZ. The popularity of using wildlife cameras in both research and recreation, paired with the advancement of citizen science projects like iNaturalist have generated a vast amount of new data on species distributions. With this new body of information we report the range of WNC now occurs over 400 km farther north, extending north of Flagstaff, Arizona. Recent records include occurrence in ponderosa pine forest that sustain sometimes heavy winter snow – an environment vastly different from the species’ normal range. The northward expansion of this meso-carnivore invites many questions about drivers of range expansion, including climate change, mesopredator release, or simple opportunism. More research into the behavior and ecology of WNC in the northern extent of their range is needed to guide understanding and potential future management of this species, its impacts, and prediction of other such range expansions.

## Introduction

White-nosed coatis (WNC; *Nasua narica* Linnaeus 1766) are semi-arboreal meso-carnivores that range from northern South America to southwestern United States (US), with a recently expanded distribution into northern Arizona. A member of Procyonidae, WNC are the only coati species found in North America (Hoffmeister 1986, Cuaron et al. 2016). Originally described by Linneaus (1766) as *N. narica*, Handley (1966) reported that Linnaeus’ description was of a specimen with a unicolored tail, while Linnaeus’ *N. nasua* had a black and white banded tail. Hoffmeister (1986) reported that the Arizona *N. nasua* were likely the subspecies *N. n. molaris* Merriam, but Hershkovits (1951) considered that subspecies to be synonymous with *N. n. pallida* Allen. A more detailed taxonomic treatment remains outstanding.

The earliest record of WNC in the US was in 1892 at Fort Huachuca, just north of the US-Mexico border (∼31°19’57”N; Wallmo and Gallizioli 1954). At that time, it was considered a rare sighting. WNC were established in the Chiricahua and Huachuca Mountain ranges during the early 20^th^ century (Cahalane 1939). However, as extensive mining and settlement took place across southeastern Arizona, sightings of WNC were still considered rare, suggesting at least that WNCs were rarely seen (Wallmo and Gallizioli 1954). A deputy game warden of the Chiricahua Mountains, Bill Lee, called WNC “infrequent” at that time, referring to it as the “Mexican cholugo” (Cahalane 1939).

By 1940, breeding WNC populations were well-described in southeastern AZ. Taber (1940) wrote that a “northern invasion” of WNC was occurring, and in 1934 WNC was added to the US mammals list because it was known to occur “sporadically on the American side of the Mexican border” (Taylor 1934, Taber 1940). Thereafter, bands of WNC began being commonly noted in the Chiricahua, Huachuca, Patagonia, and Tumacacori mountains (latitudes ∼= 31°29’20.72”; Cahalane 1939, Wallmo and Gallizioli 1954, Hall and Kelson 1959). By 1954, the local community of the Huachuca Mountains began to view WNC as a nuisance species because of injuries to dogs, depredation on chickens, and foraging on orchard fruit (Wallmo and Gallizioli 1954).

Since then, the range of WNC has continued to expand northward. More bands have become established at northerly latitudes, such as the San Carlos Indian Reservation (Kaufmann et al. 1976). Hoffmeister (1986) reported WNC from Walnut Canyon National Monument and Petrified Forest National Park; however, there are few further publications recognizing this expansion. In 2008, the International Union for Conservation of Nature and Natural Resources (IUCN) listed WNC habitat as following the oak woodlands of the southeastern region of the United States, with WNC mainly in the Chiricahua and Huachuca Mountains (∼31°33’18.58”N) because of their affinity with “hardwood riparian canyons over 1,400 m” (Samudio et al. 2008). In 2016, the updated IUCN range of WNC was drawn north of Interstate 40 in Flagstaff, AZ (∼35°11’51.93”N), showing an expansion from their records in 2008 to 2016 (Cuaron et al. 2016).

Today, WNCs have been consistently recorded in and north of Flagstaff, Arizona, a city at a longitude of ∼35°11’51.94”N (about 400 km north of the US-Mexico border) and at an elevation of 2100 m, a substantial climate difference from their primary range (Cuaron et al. 2016). These reports of coati north of the southern margin of the Colorado Plateau stimulated our inquiry into historical and recent records of their occurrence and changes in distribution. We conducted a review of databases, literature, and museum specimens to document range changes for WCN. Our records illuminate this species expansion into Arizona over the past century, as well as confirming the northernmost distribution of the species.

## Methods

To track occurrence records across time for WNC we collated data from 6 different sources: iNaturalist, peer-reviewed literature, Arizona Game and Fish Department’s (AZGF) Heritage Data Management System (HDSM) records, Museum of Northern Arizona collections, personal communications, and opportunistic camera trap photographs. For all records we collected the year of sighting, latitude, longitude, individuals sighted (i.e., band or a single individual), and photographic proof of presence. Only records with complete information were included. When roadkill or WNC tracks were recorded, we categorized these as a single individual. iNaturalist is a citizen scientist project that encourages citizens with a cell phone to photograph and document species around the world. These web-based platforms are commonly used as a tool to document species distribution, phenology, and seasonal events, among many other observations. Because records are georeferenced, have attached photographs, and are reviewed by professionals to earn the title of ‘research grade’, their authenticity is confirmed. We downloaded 322 research grade iNaturalist results in March 2021 with the parameters of “*Nasua narica”* and the filter of “Arizona” were applied (GBIF 2021). The timing of records ranged from 1999 to 2021.

To gain historical and peer-reviewed context of coati presence in Arizona we used Google Scholar and the search terms “*Nasua narica*” and “Arizona” to explore the literature. Over 700 results were returned; however, fewer than 50 mentioned WNC sightings in the state of Arizona, and only 38 of them were unique records. Publications and accounts dated from early 1892 to October 2019. When specific coordinates were not provided, an estimated geographic reference from the paper was used. If an account was repeated in the literature, only the initial publication was recorded.

We secured Heritage Data Management System (HMS) records of confirmed WNC sightings by AZGF and US Fish and Wildlife Service biologists from 1983 to 2018. Additionally, wildlife camera trap photos with geographic coordinates from the Verde were obtained from wildlife biologists for photographic evidence of WNC. Lastly, several personal communications from reliable biologists were included of WNC sightings in northern Arizona.

## Results

We documented 626 records of coati sightings in Arizona from 1892 to 2021, including: 322 from iNaturalist; 242 from HDMS reports; 39 from peer-reviewed literature (Table 1); 16 from wildlife cameras; 4 Museum of Northern Arizona specimens; and 3 from personal communications. The Museum of Northern Arizona specimens expanded our WNC distribution and history data. An adult male WNC specimen (MNA Z9-698) was collected on the Babbitt Brothers Spur Ranch at the edge of Anderson Mesa near Flagstaff, at an elevation of 1825 m on 30 January 1955. An adult male WNC specimen was recovered as a salvaged roadkill 29 km NW of Flagstaff on Highway 180 at 35°20’46.86”N on 8 July 2016. At 2,437 m, this specimen appears to be the highest elevation reported for WNC in Arizona. This specimen is curated at the Museum of Northern Arizona as MNA Z9-5512. Additional observations and photographs of WNC and their tracks were taken along the Rio de Flag Canyon at MNA, and at Walnut Canyon National Monument during the winter of 2015 and 2016.

**Table 1:** Records from literature of white-nosed coati *Nasua narica* in Arizona, United States of America.

| Year | Latitude (N) | Longitude (W) | # Individuals | Data Type | Citation |
| --- | --- | --- | --- | --- | --- |
| 1892 | 31° 33' 18.576" | - 110° 20' 58.848" | 1 | Photo | Wallmo & Gallizioli 1954 |
| 1908 | 32° 22' 29.9532" | - 109° 3' 24.2172" | 1 | Testimony | Cahalane 1939 (Bailey 1931) |
| 1914 | 31° 24' 56.2608" | - 110° 43' 48.7812" | 1 | Photo | Wallmo & Gallizioli 1954 |
| 1924 | 31° 29' 20.724" | - 110° 24' 32.2992" | 1 | Photo | Taylor, 1934 |
| 1932 | 31° 38' 39.0264" | - 109° 10' 16.6872" | 1 | Photo | Cahalane 1939 |
| 1934 | 33° 29' 53.7216" | - 111° 27' 53.4744" | 1 | Testimony | Taylor 1934 |
| 1939 | 31° 27' 42.9408" | - 111° 14' 13.254" | band | Testimony | Taber 1940 |
| 1940 | 33° 38' 14.9784" | - 109° 19' 49.4004" | 1 | Testimony | Taber 1940; Taylor 1934 |
| 1942 | 32° 5' 15.6732" | - 112° 54' 24.3756" | 1 | Testimony | Kaufmann et al. 1976 |
| 1949 | 31° 29' 20.724" | - 110° 24' 32.2992" | band | Testimony | Wallmo & Gallizioli 1954 |
| 1950 | 31° 21' 2.8764" | - 110° 16' 51.6864" | band | Testimony | Risser 1963 |
| 1953 | 31° 27' 43.4412" | - 110° 58' 53.58" | band | Testimony | Fred Fendig, Risser 1963 |
| 1954 | 32° 42' 5.1372" | - 109° 52' 17.6916" | 1 | Testimony | Wallmo & Gallizioli 1954 |
| 1955 | 31° 22' 59.0232" | - 110° 35' 7.7136" | band | Testimony | Carl Yeager, Risser 1963 |
| 1960 | 31° 27' 54.1872" | - 110° 17' 5.3484" | band | Testimony | William Brown, Risser 1963 |
| 1960 | 31° 30' 57.456" | - 110° 33' 32.922" | band | Testimony | Don Havluk, Risser 1963 |
| 1960 | 31° 25' 59.3508" | - 111° 8' 52.386" | 1 | Testimony | Risser 1963 |
| 1960 | 31° 29' 46.7196" | - 110° 51' 44.352" | band | Testimony | Charles Morgan, Risser 1963 |
| 1961 | 31° 29' 20.724" | - 110° 24' 32.2992" | band | Testimony | Sewell Goodwin, Risser 1963 |
| 1961 | 31° 21' 2.8764" | - 110° 16' 51.6864" | 1 | Photo | Risser 1963 |
| 1961 | 31° 25' 37.3476" | - 110° 17' 28.0428" | band | Testimony | James Holt, Risser 1963 |
| 1962 | 31° 33' 18.576" | - 110° 20' 58.848" | band | Testimony | Pratt 1962 |
| 1962 | 31° 39' 24.84" | - 110° 52' 9.5952" | band | Photo | Risser 1963 |
| 1962 | 31° 27' 43.4412" | - 110° 58' 53.58" | band | Photo | Risser 1963 |
| 1970 | 32° 35' 30.4764" | - 109° 50' 59.5428" | band | Testimony | Kaufmann et al. 1976 |
| 1973 | 35° 10' 18.6888" | - 111° 30' 34.4736" | 1 | Testimony | Salomonson 1973 |
| 1974 | 32° 50' 45.3948" | - 110° 38' 11.4936" | 1 | Testimony | Kaufmann et al. 1976 |
| 1974 | 32° 1' 21.432" | - 109° 21' 51.7896" | band | Photo | Lanning 1976 |
| 1975 | 32° 50' 45.3948" | - 110° 38' 11.4936" | band | Testimony | Kaufmann et al. 1976 |
| 1976 | 31° 47' 13.6428" | - 111° 34' 56.6292" | band | Photo | Kaufmann et al. 1976 |
| 1976 | 33° 55' 3.18" | - 109° 28' 32.1744" | band | Testimony | Kaufmann et al. 1976 |
| 1976 | 33° 54' 38.502" | - 109° 35' 7.5732" | 1 | Testimony | Kaufmann et al. 1976 |
| 2000 | 31° 34' 59.9988" | - 110° 25' 0.0012" | band | Photo | Hass 2002 |
| 2000 | 31° 34' 59.9988" | - 110° 25' 59.9988" | band | Photo | Hass and Valenzuela 2002 |
| 2004 | 32° 1' 21.432" | - 109° 21' 51.7896" | band | Photo | McColgin et al. 2017 |
| 2016 | North of | I-40 | 1 | Testimony | Cuaron et al. 2016 |

Most of the literature records in the early 1900s were of individuals (e.g., Taylor 1934, Cahalane 1939, Wallmo and Gallizioli 1954), suggesting that only dispersing males were moving northward into new territories at that time (Table 1, Fig. 1). However, by the mid-1900s sightings of bands became more prevalent, although those records remained closer to the US-Mexico border (Table 1). Locations used from iNaturalist confirmed observations farther north than historically recorded (Fig. 1). In 2015, WNC records at latitudes north of Phoenix began appearing more commonly (Fig. 2). All northern sightings were of individuals, until 2018 when records of bands were reported south of Sedona, Arizona at a latitude of 34°43’35.98”N (GBIF 2021). In 2014, a research project along the Verde River (latitude: ∼ 34°2’36.78”N) recovered numerous camera trap photos of WNC, primarily in sites along river canyons (Fig. 3).

**Fig. 1.**
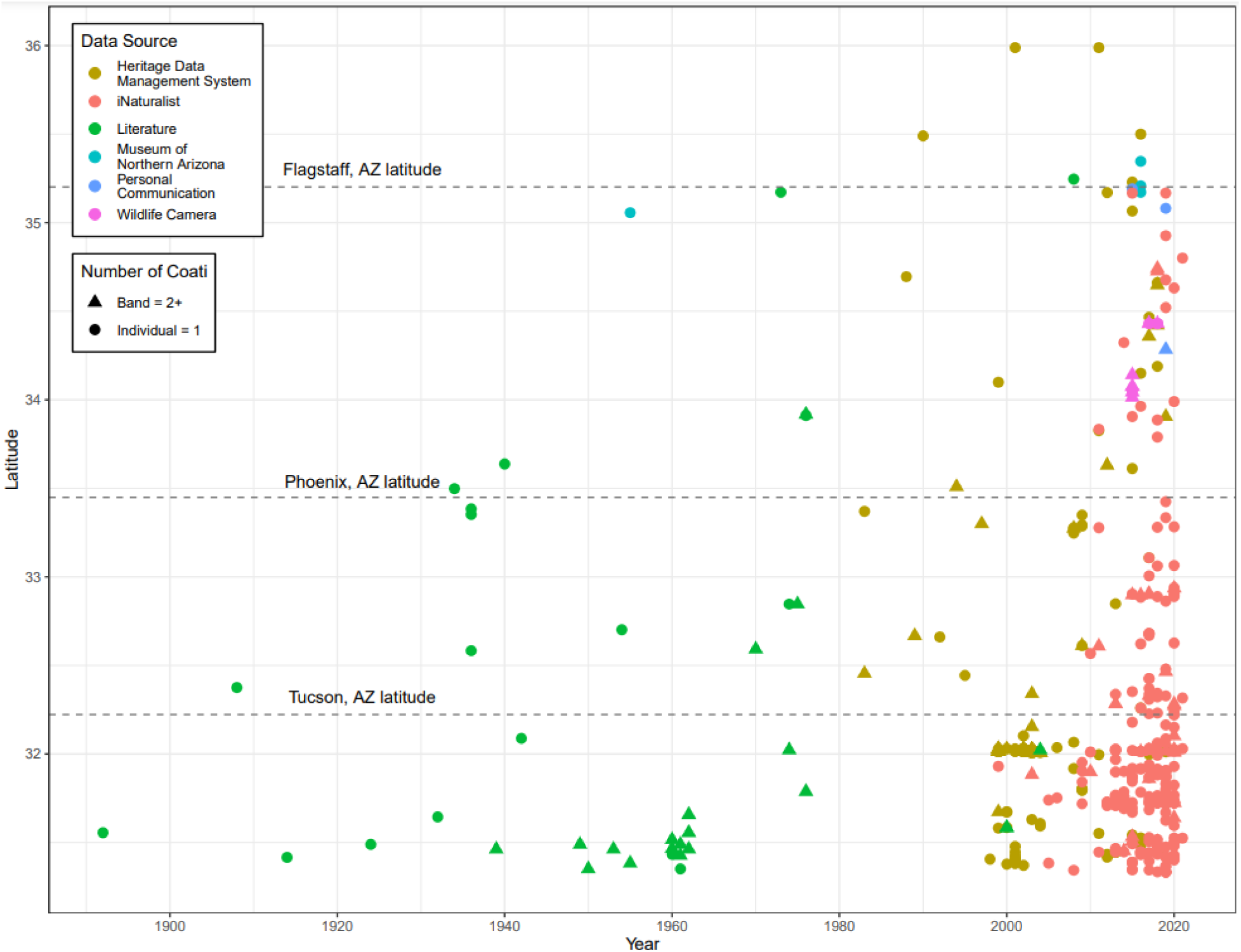
Latitudinal expansion of white-nosed coati by year (1892 to 2021). Each data point indicates either an individual or a band of coatis from one of six data sources: Heritage Data Management System, iNaturalist, literature, Museum of Northern Arizona, personal communications, and wildlife camera photos.

**Fig. 2.**
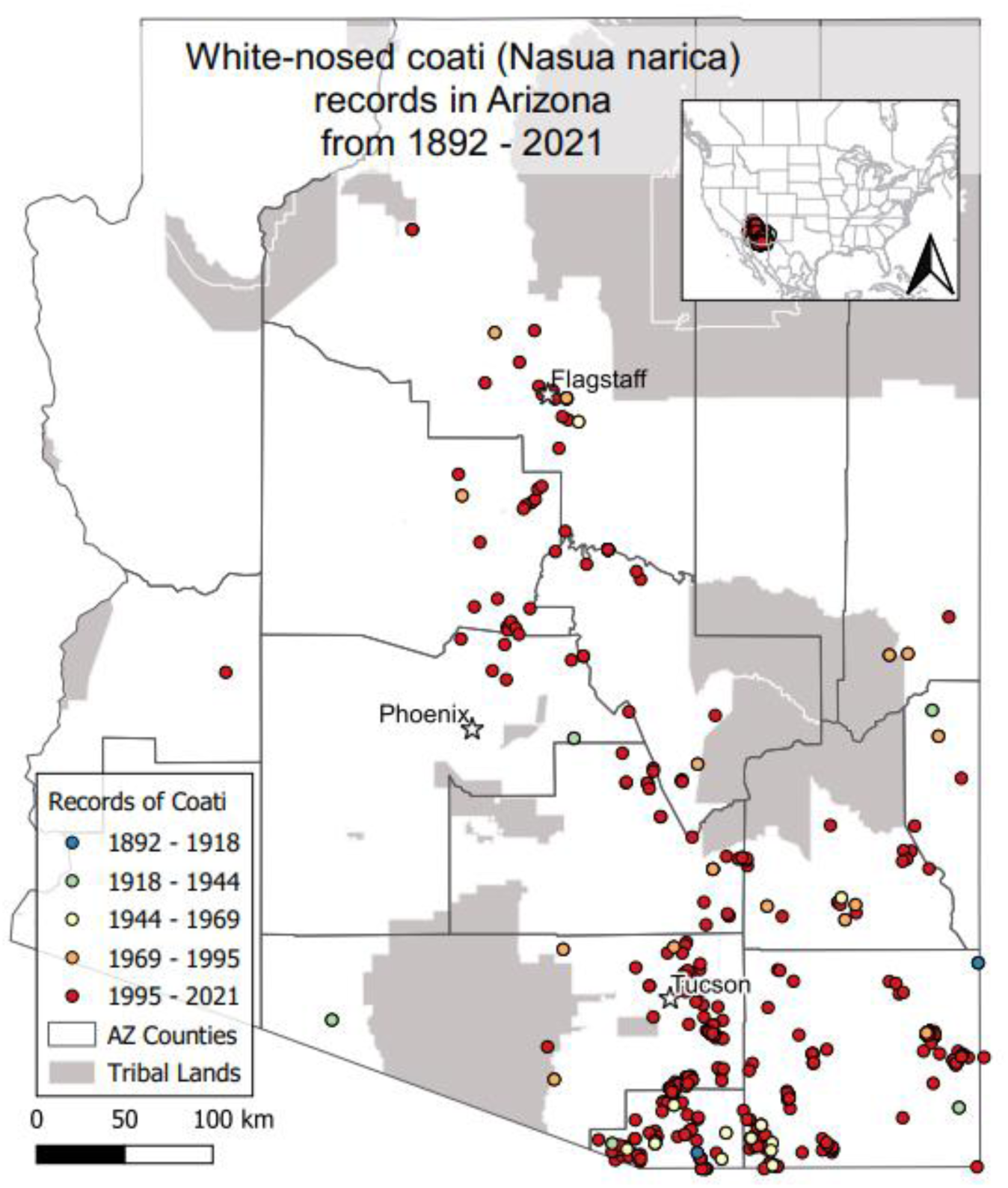
Coordinates of white-nosed coati in Arizona from 1892 – 2021. Each data point indicates a documented sighting of one or more white-nosed coatis in Arizona based on records from literature, iNaturalist, Heritage Data Management System, Museum of Northern Arizona, personal communications, and wildlife camera photos.

**Fig. 3.**
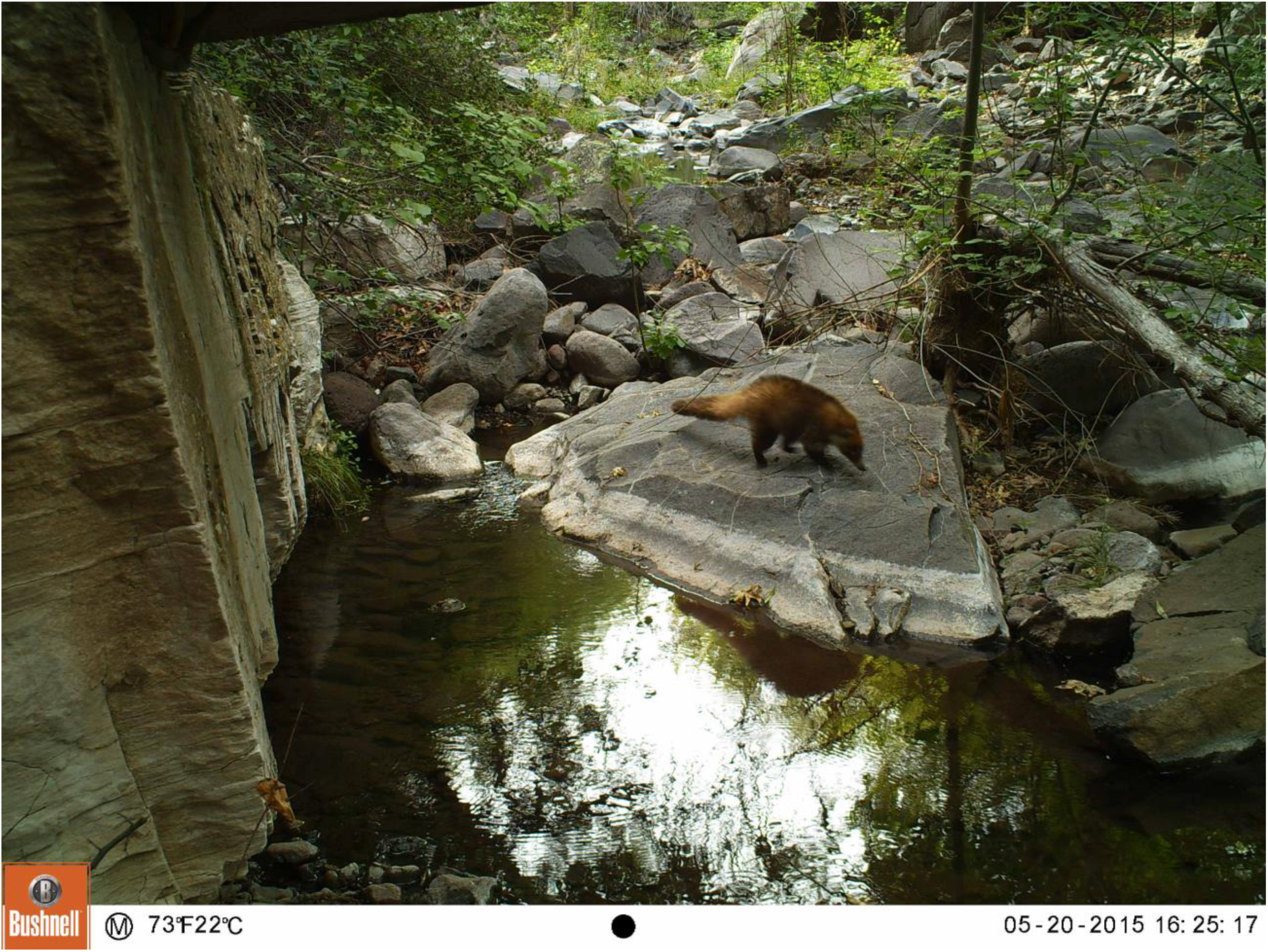
Wildlife camera photo of an individual white-nosed coati along the Verde River, Arizona in 2015.

Additionally, photographs of WNC along the Mogollon rim in Payson, Arizona were documented in 2017 and 2018 (Fig. 4). We received verified photos of an individual WNC from citizen scientists at two locations near houses within the city of Flagstaff (latitudes: 35°13’45.98” N and 35°13’17.47”N). Lastly, in Fall of 2019, two additional WNC sightings were reported by biologists in northern Arizona: one at Bear Lake Campground (latitude: 34°17’2.51”N) and the other near upper Lake Mary (latitude: 35°4’51.67”N) (Haight personal communication 2019, Tellez personal communication 2019, respectively).

**Fig. 4.**
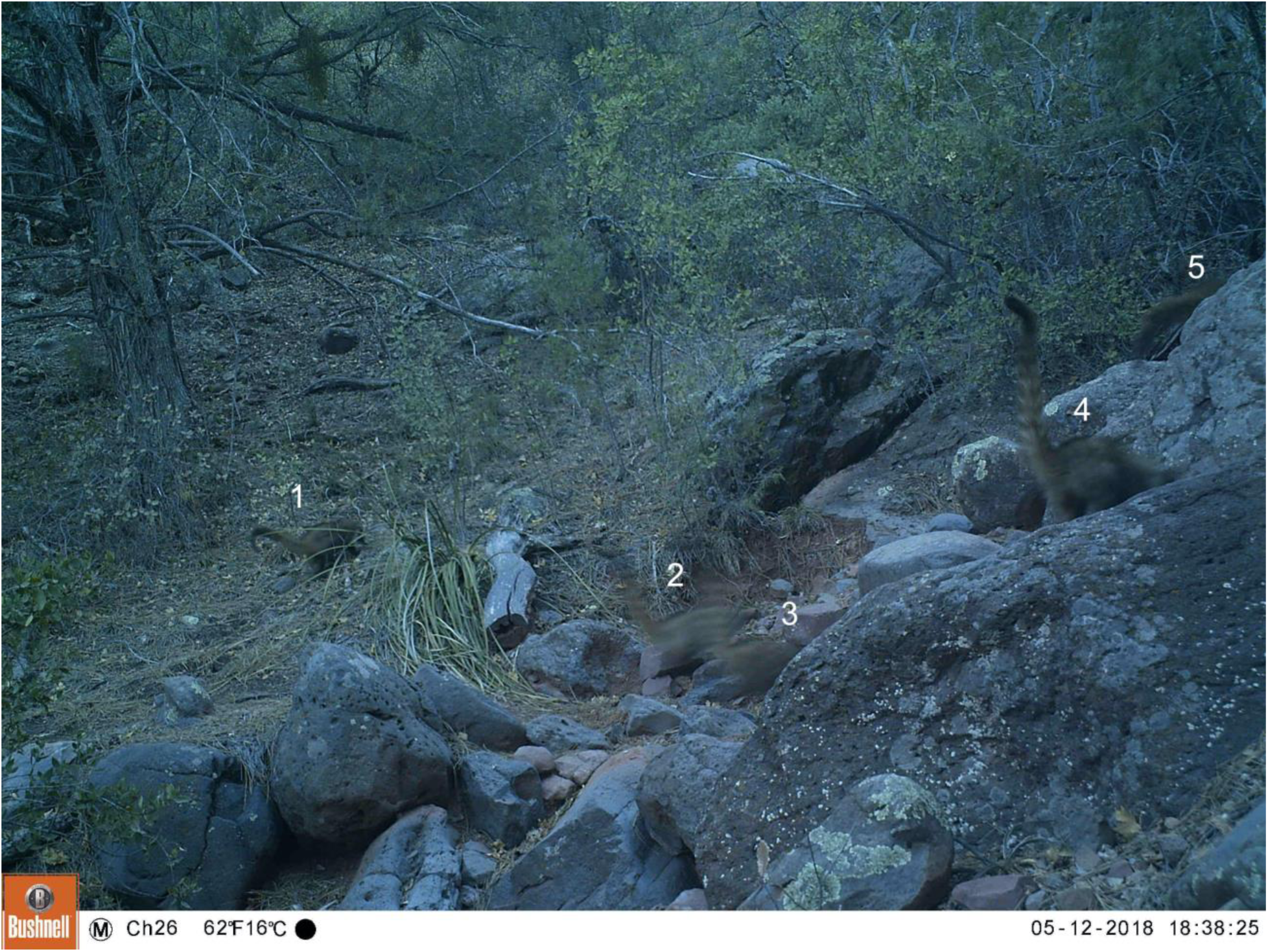
A band of 5 white-nosed coatis along the Highline Trail in Payson, Arizona in 2018.

## Discussion

It comes as no surprise that WNC, a versatile, highly adaptable, social omnivore, whose males disperse and are solitary between breeding seasons, is readily capable of colonizing new territory. Additionally, the geography of central Arizona lends itself well as a conduit for northern expansion, as hardwood river canyons connect higher and lower elevations through riparian habitat corridors (Frey et al. 2013). Our data indicate that WNC are now occurring farther north more frequently.

The opportunistic ecology and diet of WNC have likely facilitated their ability to expand into new regions. WNC occur in bands averaging 12 individuals, typically consisting of females with their young, while males are solitary or join a small bachelor troop (Smythe 1970, Hass and Valenzuela 2002). Bachelor males only interact with females during the mating season in April, and young are born in June (Smythe 1970, Hoffmeister 1986, Trovoti et al. 2010, Cuaron et al. 2016, Emmons and Helgen 2016). Outside of the mating season, adult WNC males disperse to find new territories, which enables range extension. Additionally, WNC are omnivorous, consuming insects, fruits, seeds, and forest litter invertebrates, and occasionally small-to medium-sized vertebrates (Wallmo and Gallizioli 1954, Smythe 1970, Kaufmann et al. 1976, Gompper 1995, Valenzuela 1998). It is probable that this expansive diet has facilitated their ability to exist in different habitats, adapting to local food sources as resource conditions allow (Kaufmann et al. 1976, McColgin et al. 2003, McColgin et al. 2018).

WNC have high affinity to riparian areas, especially in dry regions, which provide discrete corridors for their expansion into northern regions (Hoffmesiter 1986; Frey et al. 2013). While they have been documented in a wide array of habitats, WNC most often occupy stands of pine forest (*Pinus*), oak-pine woodland (*Quercus*), chaparral, sycamore (*Plantanus*), walnut (*Juglans*), maple (*Acer*), and other coniferous forests between 1,400 and 2,450 m elevation (Wallmo and Gallizioli 1954, Kaufmann et al. 1976, Frey et al 2013). Riparian areas provide diverse dietary and habitat needs, including: tree canopy cover, presence of water, availability of food resources, and seasonal dispersal, particularly in more arid river corridors, like the Verde River and its many tributaries (Hass 2002, Valenzuela and Ceballos 2000; Frey et al. 2013).

WNC have been recorded in Organ Pipe Cactus National Monument at what has been described as their “lowest, driest, and hottest” reported location, with only 186 mm/yr of precipitation (Kaufmann et al. 1976).

WNC are one of many Mexican-Neotropical species moving northwards as climate changes, including invertebrates and other vertebrates (Brown and Davis 1995). For example, all butterfly species added to the Grand Canyon National Park list in the past 70 years, and which are native to North America, have been Mexican-Neotropical species (Garth 1950, Stevens 2012). A similar case may be that of the Central American Masked Clubskimmer dragonfly *Brechmorhoga pertinax*, which has been reported breeding in northern Arizona (Stevens and Bailowitz 2009). Several bird species native to North America also are undergoing northward range expansion, such as Great-tailed Grackle *Quiscalus mexicanus* (Wehtje 2003). Other non-flying mammals expanding their ranges northward in the US Southwest (in addition to coyote *Canis latrans*) include collared peccary *Pecari tajacu* and American hog-nosed skunk *Conepatus leuconotus leuconotus* (Holton et al. 2021). Peccary range expansion appears to be related to the slightly warmer winter temperatures to which the Southwest is now subject (Bender et al. 2014), and both peccary and hog-nosed skunk have been detected on the north side of the Colorado River in the Grand Canyon (Stevens 2013; Holton et al. 2021). Too few distributional data are available to understand the ranges of many bats, but the phyllostomid Mexican long-tongued bat *Choeronycteris mexicana* has been detected in Grand Canyon National Park (Stevens 2013). Lastly, jaguar *Panthera onca* have recently been documented moving between the borderlands into southeastern Arizona along the same approximate route as WNC (Culver et al. 2016). Thus, WNC appears to be one of many Mexican-Neotropical taxa expanding their ranges northward into the southwestern US.

During the last century, the Arizona landscape has changed dramatically (Hutchinson et al. 2000, Jenerette and Wu 2001, Turner et al. 2003). Humans have altered habitat conditions, allowing allowed WNC to cross some of the arid landscape gaps that previously restricted them to southern Arizona. Wildlife populations and habitat availability also have greatly changed during this interval. With fewer large carnivores and less intact riparian forest cover, mesocarnivore release may have occurred, allowing WNC to proliferate and expand their range.

Today, WNC populations in the Chiricahua and Huachuca Mountains are well-established and often attract public attention in the Chiricahua National Monument, where they are seen daily by visitors. Continued individual sightings in Flagstaff north of Interstate 40, and with bands documented at higher latitudes indicate that WNC will continue to move northward. WNC are well studied in Central America; however, data on their behavior and ecology at the northernmost extent of their range are outstanding. Although we can speculate on driving factors, we have no direct evidence to suggest why they are expanding northward now. If the current trends observed in our data continue, WNC may soon disperse to Grand Canyon National Park and the Colorado River corridor, and other suitable habitats in northern Arizona. Such changes may be of interest to land managers, who might see WNC as welcome guests or regard them as invading “southern pests”.

Dispersal and range expansion are natural processes; however, relatively rapid range expansions also can indicate larger issues that may warrant scientific and managerial attention. Further research is needed to understand WNC habitat selection and interactions with northern species. Cuaron et al. (2016) stated that large-scale habitat loss threatens US populations of WNC, and warn that US populations may be becoming genetically isolated from Mexican populations. To better understand the potential impact of WNC on other wildlife and WNC movement corridors, camera trapping, animal collaring, genetic assessments of population connectivity, and diet assessment should be conducted to develop information about their distribution, home range size, and behavior. However, in the short term it will be important to conduct research on the mechanisms driving WNC range expansion, and how their presence and occurrence in developed areas might impact existing management actions. Such data can provide the basis for a predictive habitat model to forecast WNC movement and habitat use, and to improve prediction of adaptive management options for future vertebrate communities (e.g., Frey et al. 2013). With global climate change continuing to reshape southwestern landscapes, more information on transient and rapidly adapting species like WNC and their responses to these changes is essential for understanding the future of southwestern biotic assemblages.

## ACKNOWLEDGEMENTS

We thank the Museum of Northern Arizona for supporting L. Stevens’ work, and Janet Gillette for assistance with specimen data from the MNA collection. We also thank Arizona Game and Fish’s Heritage Data Management department for their contribution of records. Lastly, we thank the field volunteers (Chelsey Tellez, Ray Tellez, Samantha Lloyd, and Sandy Leander) who helped manage and retrieve cameras and our pilot from the Arizona Pilots Association Tommy Thomason.

